# First report on Bacterial Diversity of Potable Spring water of Indian Himalayan Region

**DOI:** 10.1101/320275

**Authors:** Ashish Kumar Singh, Saurav Das, Samer Singh, Varsha Rani Gajamer, Nilu Pradhan, Yangchen D. Lepcha, Hare Krishna Tiwari

## Abstract

Water quality of a region directly corroborates with the health index of people. People in the Himalayan hills mainly depend upon the spring water for potability. To determine the microbial ecology of the spring waters of Sikkim, the variable region of 16S rRNA has been sequenced using Illumina MiSeq. Phylum wise annotation showed the East and North district are mostly dominated by *Proteobacteria* (41% and 35.80%), whereas West and South district is dominated by *Planctomycetes* (38.46%) and *Verrucomicrobia* (33%). The consistent dominance phyla in the all the four districts were *Bacteriodetes* (34-24%) which was highest dominancy in North district and lowest in wets district. Genus wise distribution showed the abundance of *Brevifolis, Flavobacterium, Verrucomicrobia subdivision3, Emticica, Cytophaga, Prosthecobacter, Planctomycetes, Varivorax, Arcicella, Isosphera, Sedimunibacterium* etc. The East district showed highest dominancy of genus *Emticicia* whereas *Planctomycetes* in the West district. The North district was mainly dominated by genus *Arcicella* and *Brevifollis* in the South district. North on the antonymous showed totally different sets of microbial diversity. North district showed an abundance of *Arcicella, Planctomycetes, Schlerensia* and *Azohydromonas*. The heat map produced by Bray Curtis distance method produced three clusters which showed the close relationship between West and East district microbiome that further related to South district. The sample of North district formed out group that showed different community structure from other three districts. The principle component analysis was showed that the east and South district samples are closely related and distantly correlated to the west Sikkim, but the North district showed completely different microbial community. The canonical correspondence analysis showed correlation between bacterial diversity and hydrochemistry and it was found that the bacterial diversity was influenced by the concentration of different metallic ions like sodium, calcium, barium and iron. This is a first report from the Eastern Himalayan region of India and it largely enhances our knowledge about the microbial structure of potable spring water of Eastern Himalayan. This study is useful for Government of India as well as the state government to adopt the different strategic treatment procedures to improve the quality of water that is supplied to the community resides in the Himalayan regions and solely dependent on this untreated spring water.

## 1.0 Introduction

Microbial communities are the key members of many ecosystems on the earth (Lagkouvardos *et al*., 2016). The word microbial communities can be described in the terms of richness or evenness as well as composition i.e. abundance of taxa and genes in the sample (Rothschild, 1991). They play a significant role in crucial biological functions ranging from nitrogen and carbon cycles in the environment to regulation of metabolic and immune responses in inhabited animal and human hosts (Rothschild, 1991; Hörmannsperger *et al*., 2012; Lagkouvardos *et al*., 2016). Hence, the study of microbial ecology is important for proper understanding of these microbial communities and their functional impact (Lagkouvardos *et al*., 2016).

Around one-third of global freshwater reserves are placed in subsurface streams and aquifers which represent as a freshwater source for human consumption as well as irrigation (Hemme *et al*., 2015). As good quality freshwater is an important natural source for domestic as well as the industrial purpose which is gradually exposed by a human (Van Rossum *et al*., 2015; Vörösmarty *et al*., 2010) as well as anthropogenic activity that affect the quality of water (Vörösmarty *et al*., 2010). Throughout the world contamination of water bodies by waterborne pathogens and disease caused by them are a major water quality concern (Pandey *et al*., 2014). The potential host-specific nature of the microbes they have long been used as an indicator of poor water quality (Staley *et al*., 2013). Although, these tests for detection of pathogens are based on culture-dependent methods which carry both false positive as well as a false negative result (McLain *et al*., 2011; Staley *et al*., 2013). The exploration of microbial communities through culture-dependent methods were used from ancient time, but on the standard laboratory media less than 1%, bacterial species of the environmental communities were culturable (Staley and Sadowsky, 2016). To overcome the traditional culture methods, now metagenomics delivers a suitable method for monitoring the environmental communities via high-throughput manner. Such techniques have revealed unprecedented taxonomic and functional diversity in aquatic and terrestrial habitats (Sogin *et al*., 2006). The term metagenomics describes as the sequencing of total DNA isolated directly from the environmental samples i.e. ‘metagenomes’ which simultaneously provides the access of genetic information of microbial communities from the mixed environment (Dinsdale *et al*., 2008). By applying this approach, functional and taxonomic microbial diversity can be described and then changes in communities can be monitored over time and space in response to anthropogenic or environmental impacts related to human health (Port *et al*., 2012). The growing accessibility of next-generation DNA sequencing (NGS) methods has greatly advanced our understanding of microbial diversity in medical and environmental science (Lee *et al*., 2017). The Next-generation sequencing (NGS) methods have prominently increased sequencing throughput by using massively parallel sequencing (Sogin *et al*., 2006). The amplification of small but variable regions of the 16s rDNA i.e. V3, V4, V5 or V6 hypervariable regions has provided very profound sequencing of bacterial ecology. This method has used for the identification of very rare populations which is present in very low abundance that may necessary for the functional diversity and ecosystem stability (Sogin *et al*., 2006; Staley *et al*., 2013). Several bacterial communities from different environmental samples like marine waters (Brown *et al*., 2009), soils (Jones *et al*., 2009), wastewater (Sanapareddy *et al*., 2009), contaminated water (Das *et al*., 2017) and several human microbiomes were successfully characterized (Staley *et al*., 2013; Sanapareddy *et al*., 2009).

The range of Himalaya is being the source of innumerable perennial rivers, streams as well as springs and the mountain peoples, are largely dependent on this spring water for their sustenance. In mountain regions, there are several natural discharges of groundwater from several aquifers in the form of Springs. The mountain Springs are locally known as Dharas which is in most cases unconfined. In Sikkim, most of the springs are denoted as ‘Devithans’ which are considered as sacred and protected as well as prevented from any living interferences to maintain the holiness of the respected springs. In Sikkim, about 80% of the rural households population depend on spring water for their drinking as well as household purpose (Tambe *et al*., 2012). In the current study, we tried to explore the overall bacterial ecology of spring water of rural as well as urban areas of Sikkim, North East India on the basis of a culture-independent study of conserved region V3-V4 of 16S rRNA gene. Along with microbial ecology physicochemical analysis was also done to correlate with the diversity study.

## 2.0 Materials and Methods

### 2.1 Site Description

The northeastern region of India is one of the biodiversity hotspots in the country. It consists of eight states which are often referred to as seven sisters (Assam, Arunachal Pradesh, Manipur, Nagaland, Tripura, Meghalaya, Mizoram) and one brother state (Sikkim). Sikkim is the second smallest State of India lying in ecological hotspot region of the lower eastern Himalayan belt. It has both alpine and subtropical climate with a high mountain range of widespread altitude variation (300-8598 m). It is the host to Kanchenjunga (8598m), the highest peak in India and third highest on earth. The state is subdivided into four districts – East Sikkim, West Sikkim, South Sikkim and North Sikkim. This mountain state is a wonder when it comes to water. It is a house of waterfalls, springs, river, and lakes, which are always an attractant to tourists and nature lovers. Rainfall is the primary mode of recharge for most of the surface water sources. The precipitation of the water in the mountain range causes surface run-off which often takes the shape of streams, springs, and kholas (local name for springs). These springs remain as an important source of potable water for the local residents **(Fig. 1)**.

**Fig. 1:**
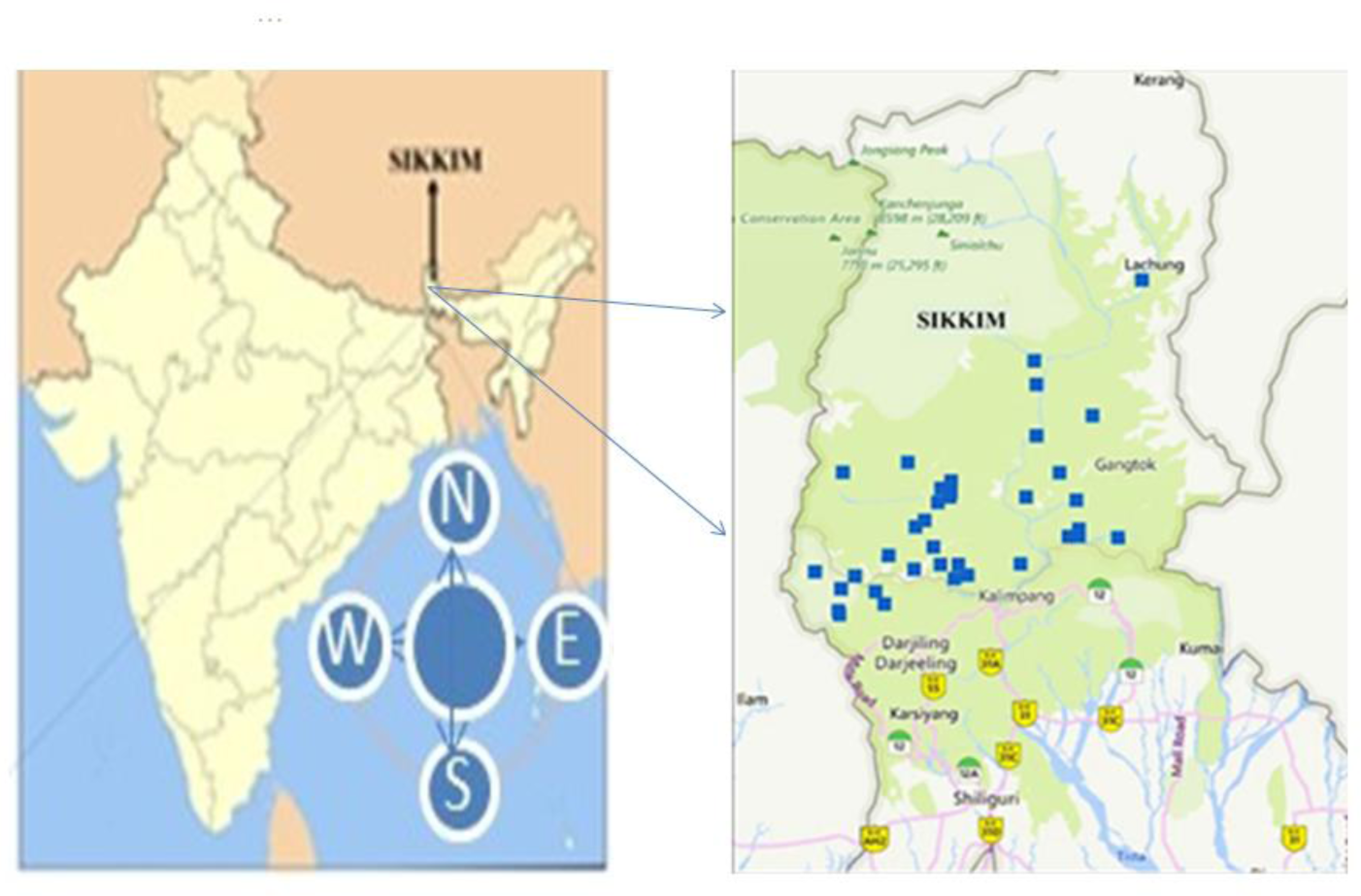
Map of Study are with regional reference with respect to Map of India. The location is marked with respect to their GPS location [lattitude and longitude] (Supplementary File – S1: Name of the place with their GPS location).

### 2.2 Sampling Procedures and Processing

A total of 40 samples, 10 from each district was collected in nalgene wide moth sterile sample bottle (ThermoFisher Scientific, USA) for microbiome analysis. Primary physical parameter including pH and temperature were tested using portable multimeter probe(Hi-media, Mumbai). The physicochemical parameters were analyzed with Induced Coupled Plasma-Mass Spectrometry (ICP-MS) (**Table. 1**). For total microbial diversity analysis, collected samples were thoroughly mixed with the equal ratio (1:1) based on the distribution like East, West, South and North district separately on different vials to prepare four composite samples of 100ml, representative of the four districts.

### 2.3 DNA Extraction

DNA from water sample was extracted using DNeasy PowerWater Kit (MO BIO Laboratories, Carlsbad, CA, USA) in accordance with the manufacturer’s instruction. Quality of the DNA was checked on 0.8% agarose gel and DNA was quantified using Qubit Fluorometer (Vr. 4.0; Thermofisher Scientific, USA), which has a detection limit of 10 - 100 ng/ μl (**Table. 1S**).

### 2.4 Metagenomic Sequencing

#### 2.4.1 Preparation of 2x300 MiSeq library

The amplicon libraries were prepared using Nextera XT Index Kit (Illumina inc.) according to 16S metagenomic sequencing library preparation protocol (Part # 15044223 Rev. B). For amplification of eubacterial and archaeal V3-V4 region, specific primer pair V3 Forward: GCCTACGGNGGCWGCAG and V4 Reverse: ACTACHVGGGTATCTAATCC were synthesized and used. The amplified product was confirmed on 1.2% agarose gel. The adaptor-ligated amplicons were amplified using i5 and i7 primer for the addition of multiplexing index sequence required for cluster generation (P5 and P7 standard of Illumina cycle sequencing). The amplicon library was purified with AMpureXP beads (Beckman Coulter Genomics, Danvers, MA, USA). The amplified library was validated by Bioanalyzer 2100 (Agilent Technologies) using High Sensitivity (HS) DNA chips and concentration was quantified by Qubit fluorometer **(Table. S2)**.

#### 2.4.2 Next-generation Sequencing and Sequence Analysis

Based on the data as obtained from the Qubit fluorometer and the bioanalyzer, 500µl of the 10 pM library was loaded into MiSeq cartridge for cluster generation and sequencing using Paired-end sequencing method. The sequence data were analyzed using the QIIME (Quantitative Insights Into Microbial Ecology) version 1.8.0 software program (Wang *et al*., 2018). The adapter trimmed sequence was subjected to pre-processing for De-replication, Singleton removal, Chimera filtering with SolexaQA. Sequences with Phred score lower than 20, ambiguous bases having primer mismatch and low read length less than 100bp were removed. Sequences were grouped into Operational Taxonomic Units (OTUs) at 97% sequence similarity level using UCLUST (Edgar, 2010) and a consensus taxonomic classification was assigned to each representative OTU using the UCLUST classifier with a Greengenes 13.8 reference database (DeSantis et al., 2006).

### 2.5 Statistical Analysis

Normalization of the OTUs relative abundance data was performed by log transformation log10 (xi+1). Diversity analysis of the microbial ecology of spring water was authenticated with alpha, Shannon and Simpson diversity indices and test of significant difference were done by one-way using PAST software (ver. 3.1). Principle component analysis was carried out using R statistics (package – facto extra). Heatmap was constructed using R – statistics (package – ggplot) using Bray Curtis distance method to visualize the comparative differences in the microbiome community by excluding the taxa with less than <0.1% of abundance. Canonical Correspondence Analysis (CCA) was used to determine the correlative relationship between the microbial community (Phylum level) and the physicochemical parameters (R-Statistics, Package – Vegan).

### 2.6 Data Availability

The sequence obtained through high throughput sequencing method was submitted to NCBI Server and available under the accession number of East Sikkim Fresh water is SRX4016322, West Sikkim is SRX4016323, South Sikkim SRX4016320 and North Sikkim is SRX4016321.

## 3.0 Results

### 3.1 Physicochemical Properties of the Samples

Different physicochemical parameters were determined by multiprobe parameter (on site) and IC-PMS as mentioned in **Table. 1**. All the water samples collected were normal to alkaline in nature. The temperature was 22 – 25°C in most of the samples except those collected from the North Sikkim, which showed a little drop in the overall temperature with a range of 17 – 22°C.

**Table 1:**
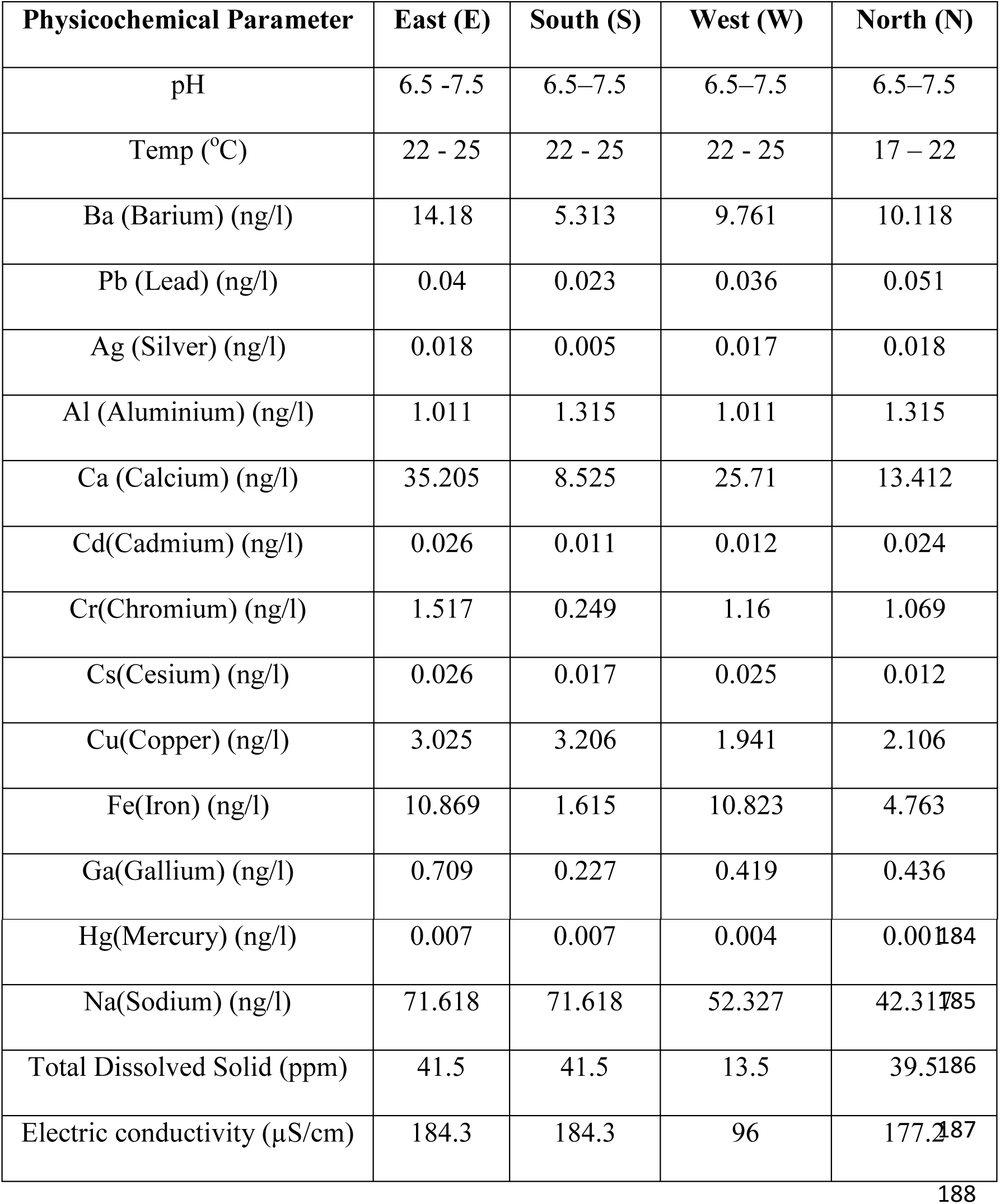
Physicochemical Parameter of the composite samples. Elemental analysis was done with ICP-MS and pH, the temperature was recorded at the site with a multiprobe parameter (Hanna Instruments).

### 3.2 Refraction Curve and Diversity Index

Rarefaction curve and diversity index showed **(Fig. 2)** most of the microbial species were sequenced and identified, and significant variations in diversity among the water samples of four districts were observed. Samples from North and East districts showed high diversity as compared to the south and west districts, with Simpson index of N: 0.6988 & E: 0.6928 and S: 0.7499 & W: 0.7356 respectively (Supplementary Table. S3).

**Fig. 2:**
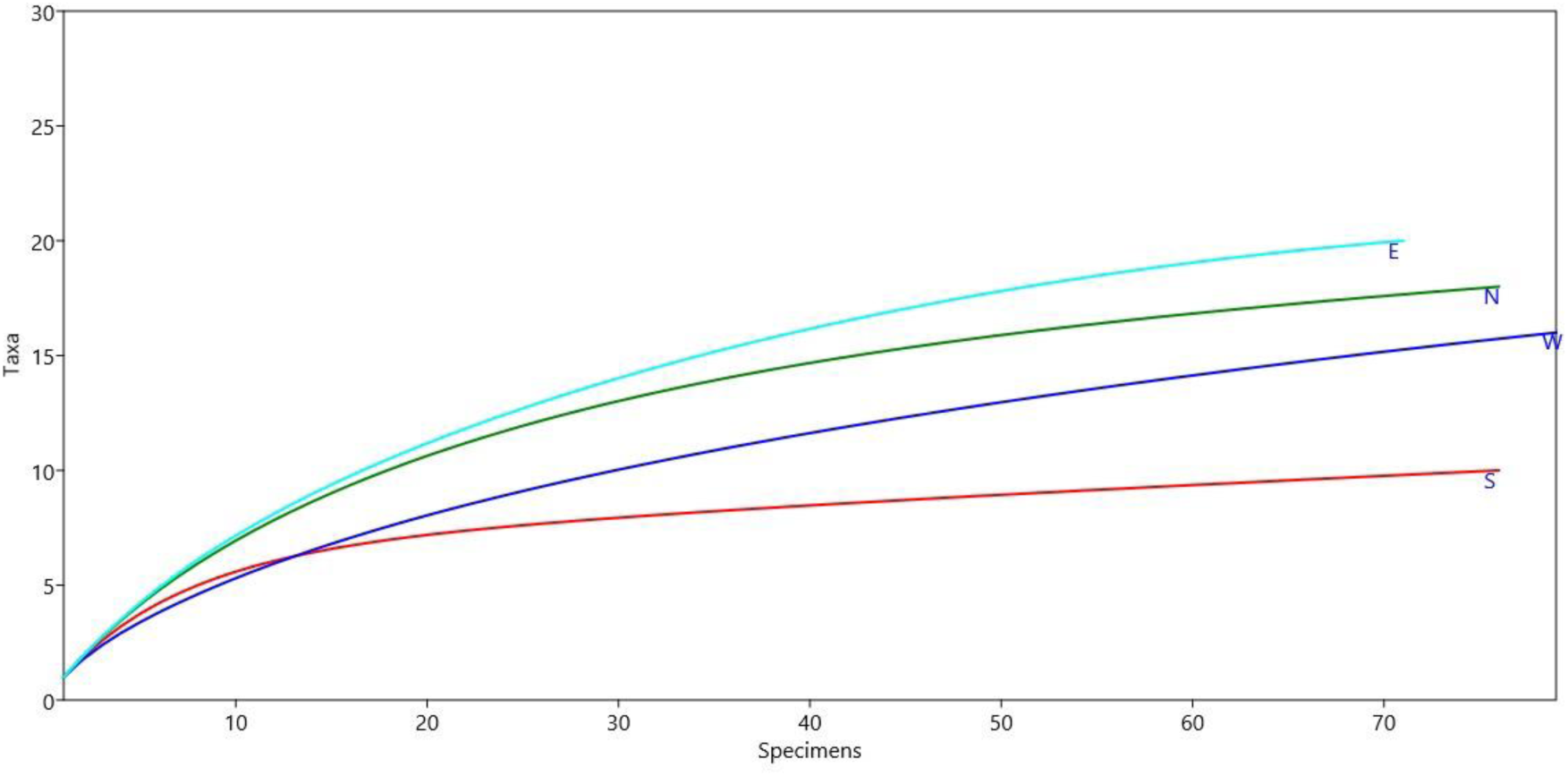
Rarefaction curve analysis of four districts of Sikkim, East (E), West (W), South (S) and North (N).

### 3.3 Microbial Diversity

The phylum wise distribution revealed the dominance of *Bacteroidetes, Proteobacteria, Verrucomicrobia, Planctomycetes, Armatimonadetes* and to lesser extent *Parcubacteria, Actinobacteria* with other minor groups like *Aquificae, Firmicutes, Hydrogenedentes, Acidobacteria, Nitrospirae, Deinoococcus-Thermus*, and *Chloroflexi* **(Fig. 3a)**. Diversity ratio was significantly different within the samples of each district. Spring water from east district showed the dominance of *Proteobacteria* (41%), Bacteroidetes (33.21%) followed by *Verrucomicrobia* (15.12%) and *Planctomycetes* (7.54%). Major phylum from the spring water of west district was *Planctomycetes* (38.46%), *Bacteroidetes* (24.04%), *Proteobacteria* (21%) and with least abundance of *Armatimonadetes* (8.08%) and *Verrucomicrobia* (6.47%). Potable spring water of south district was predominated by *Verrucomicrobia* with 33% of relative abundance followed by *Bacteroidetes* (25.46%), *Proteobacteria* (23.25%) and *Planctomycetes* (13.05%). Sample from north district showed the dominance of *Proteobacteria* (35.80%) followed by *Bacteroidetes* (34.90%) and *Planctomycetes* (22.16%). Distant from the validated phyla few *Candidatus* phyla were also recorded though at lower abundance throughout the samples *viz. TM6*, *WWE3, SR1, SBR1093, FBP, CPR2*.

**Fig. 3:**
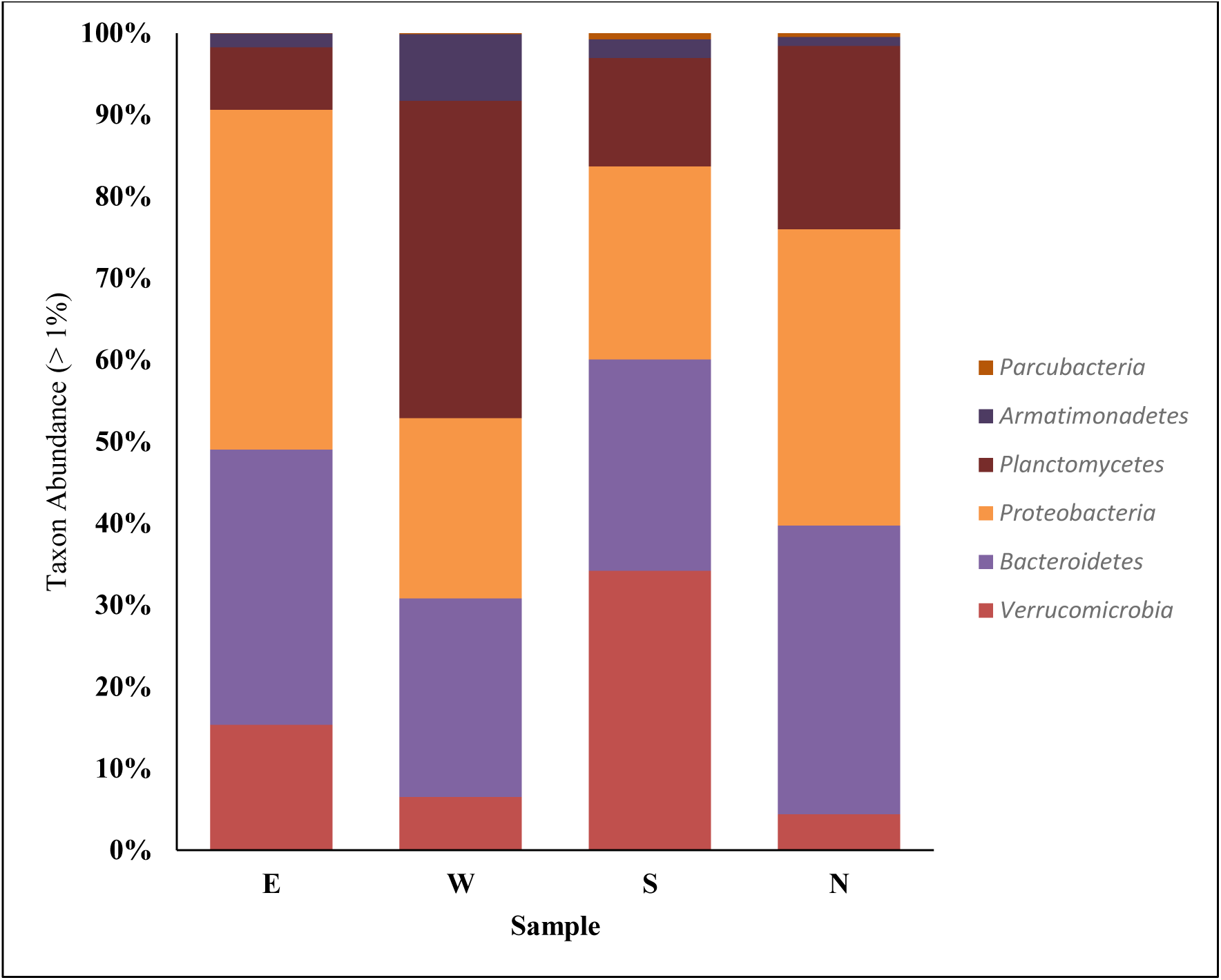
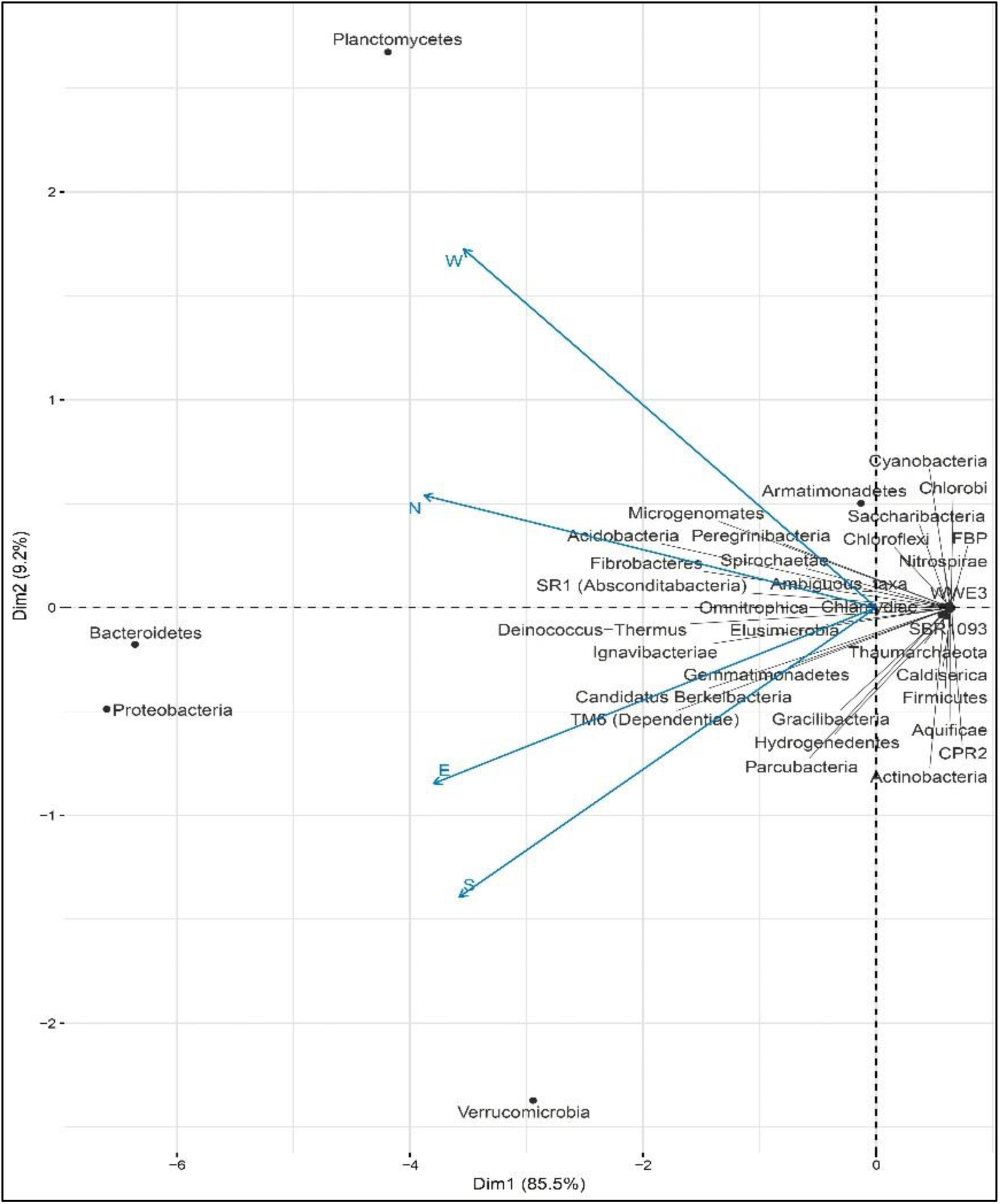
Relative abundance of phylum as identified from the spring water samples of East (E), West (W), South (S) and North (N) districts. Major phylum as found were *Bacteroidetes, Proteobacteria, Verrucomicrobia*, and *Planctomycetes* and as represented in Bar-plot **(a)** and PCA-Biplot: The principal component analysis using unweighted unifrac distance method showed the microbial distribution in the sample of west and north, east and south are correlated **(b)**.

Family wise distribution **(Fig. 4)** showed the major dominance of *Verrucomicrobiaceae, Verrucomicrobia subdivision*3*, Planctomycetaceae, Comamonadaceae, Cytophagacease, Rhodospirillaceae, Sphingomonadaceae, Chitinophagaceae, Glycomycetaceae, and Flavobacteriaceae*. Springs from the east district were mainly dominated by *Cytophagacease (15.22%)* while the west district was dominated by *Planctomycetaceae* (38.34%). South district showed the dominance of family *Verrucomicrobiaceae* (19.84%) and the north district was dominated by *Planctomycetaceae* (22.12%). Some of the major genus found in the spring water was *Brevifolis*, *Flavobacterium, Verrucomicrobioa subdivision*3*, Emticicia, Cytophaga, Prosthecobacter, Planctomycetes, Varivorax, Arcicella Isosphera, Sediminibacterium, Acinetobacter, Chitinophaga, Rhodopirellula, Tenacibaculum, Flexibacter, Ustilago, Lactobactria, Flectobacillus, Pandoraea* and *Geobacillus*. East district was dominated by genus *Emticicia* (14.83%), west district by *Planctomyces* (36.89%), south district by *Brevifollis* (19.02%) and north district by *Arcicella* (18.29%) **(Fig. 5)**.

**Fig. 4:**
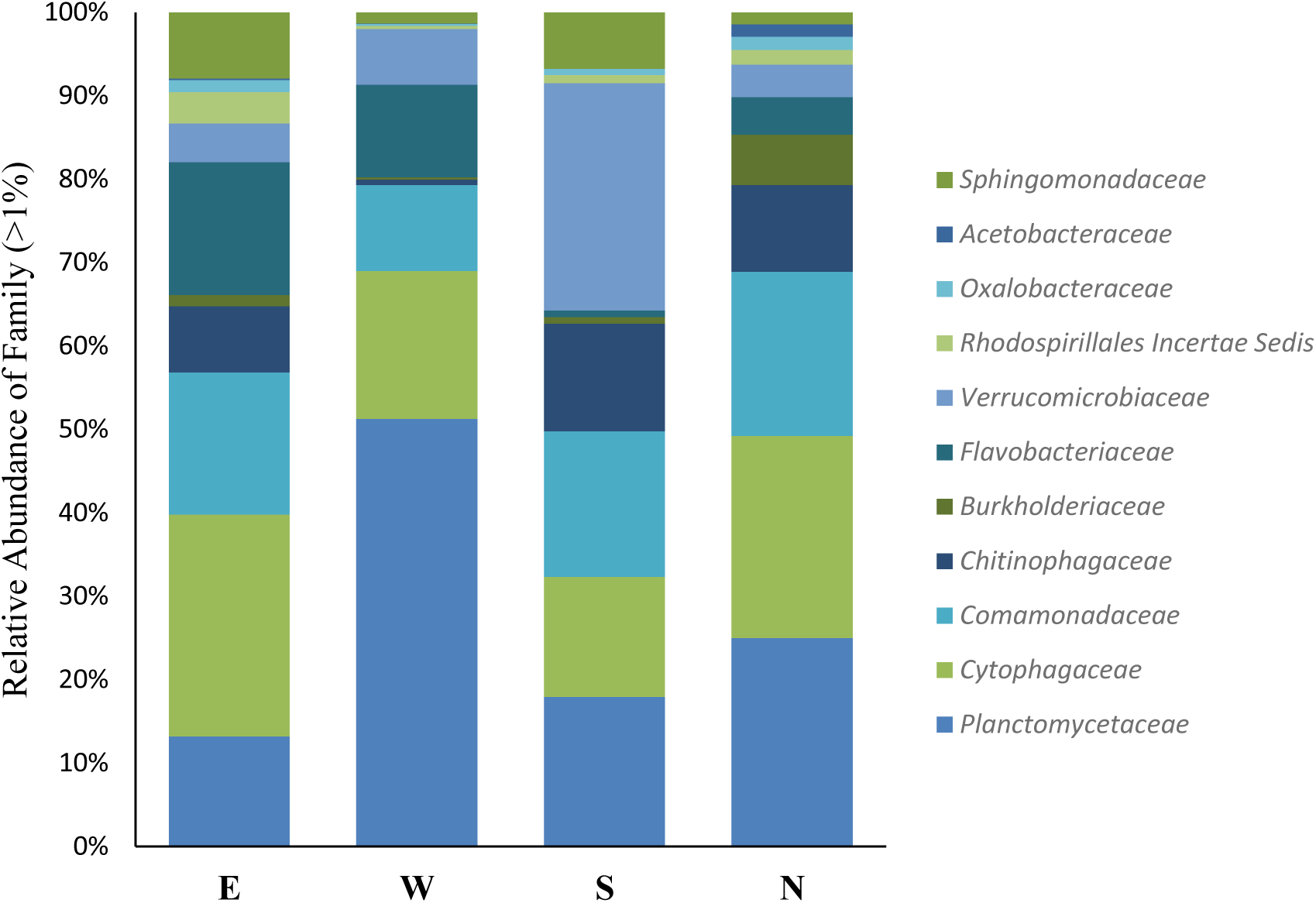
Family wise distribution of microbial flora across four districts, East (E), West (W), South(S) and North (N).

**Fig. 5:**
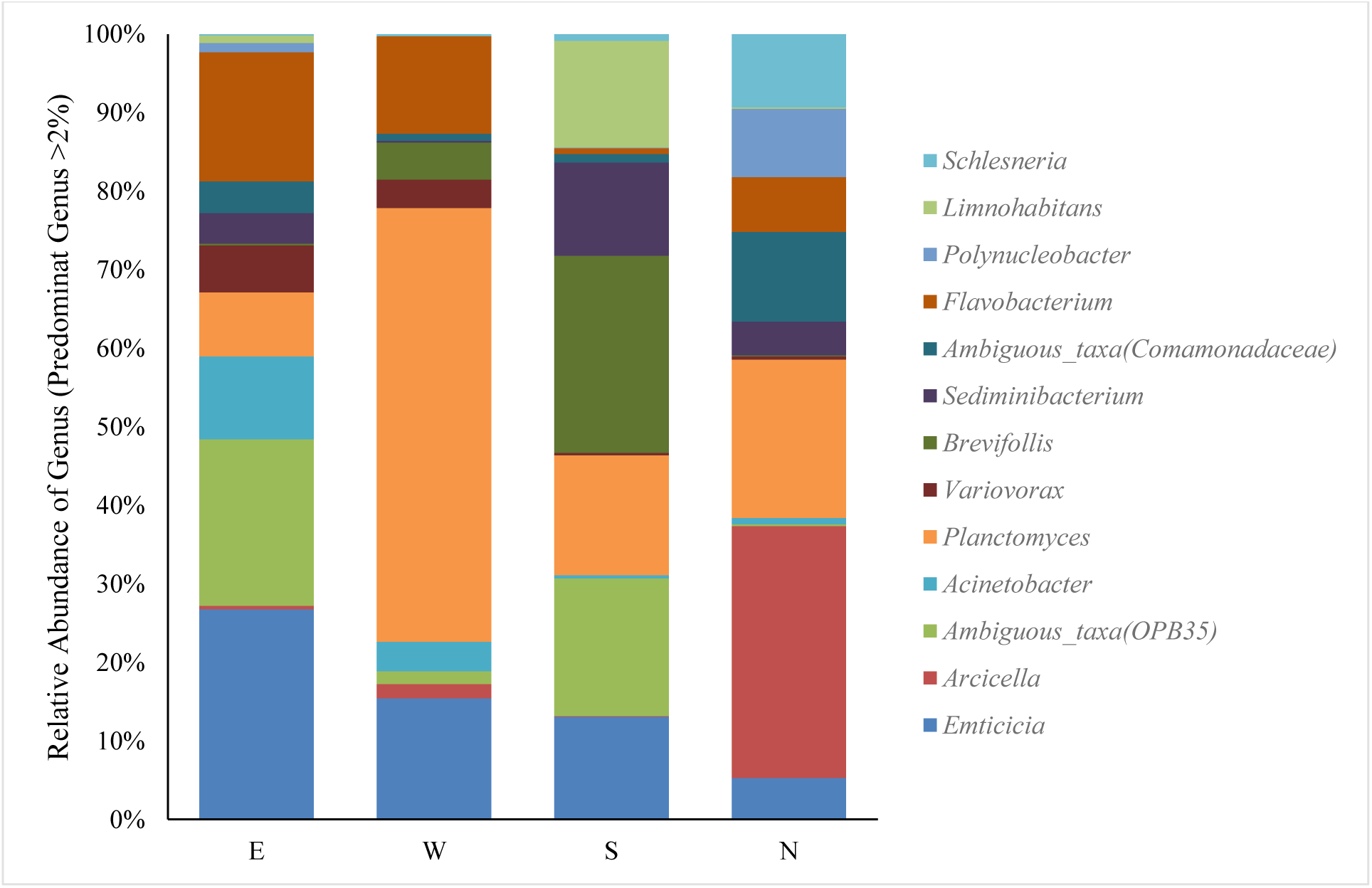
Relative Abundance of Genus in the spring water of four districts of Sikkim. The chart was prepared excluding the genus with less than 50 reads correspondingly in all the samples.

### 3.4 Comparative Community Analysis

Comparative community analysis of the Microbiome of spring water from the four districts showed a significant correlation pattern. Genus *Arciella, Planctomycetes, Polynucleobacter, Schlesneria* were mainly found in the spring water of north district with comparably higher relative abundance than the other districts. Genus *Brevifolis, Limnohabitans*, *Sediminbacterium* were abundant in the waters of south district. Except in south district, genus *Flavobacterium* was found to be predominant in the spring water of north, east and west districts. Although genus *Planctomycetes* were found in all the districts, its dominance in the West district was found to be high. *Emticicia* was a dominant genus of the East district with relatively low abundance in west and south district and in North district presence of genus *Emiticicia* was not recorded. Heatmap produced with the Bray Curtis distance method produced three clusters showing the close relationship between the Microbiome of west district and east district which is again related to the South district. North produces the out-group showing the different community structure from the rest of three districts **(Fig. 6a)**. This can also be observed from the principal component analysis where the Microbiome of east and south are showing close correlation and which is again distantly related to the Microbiome of the west but north have totally different microbial community structure (**Fig. 6b**).

**Fig. 6:**
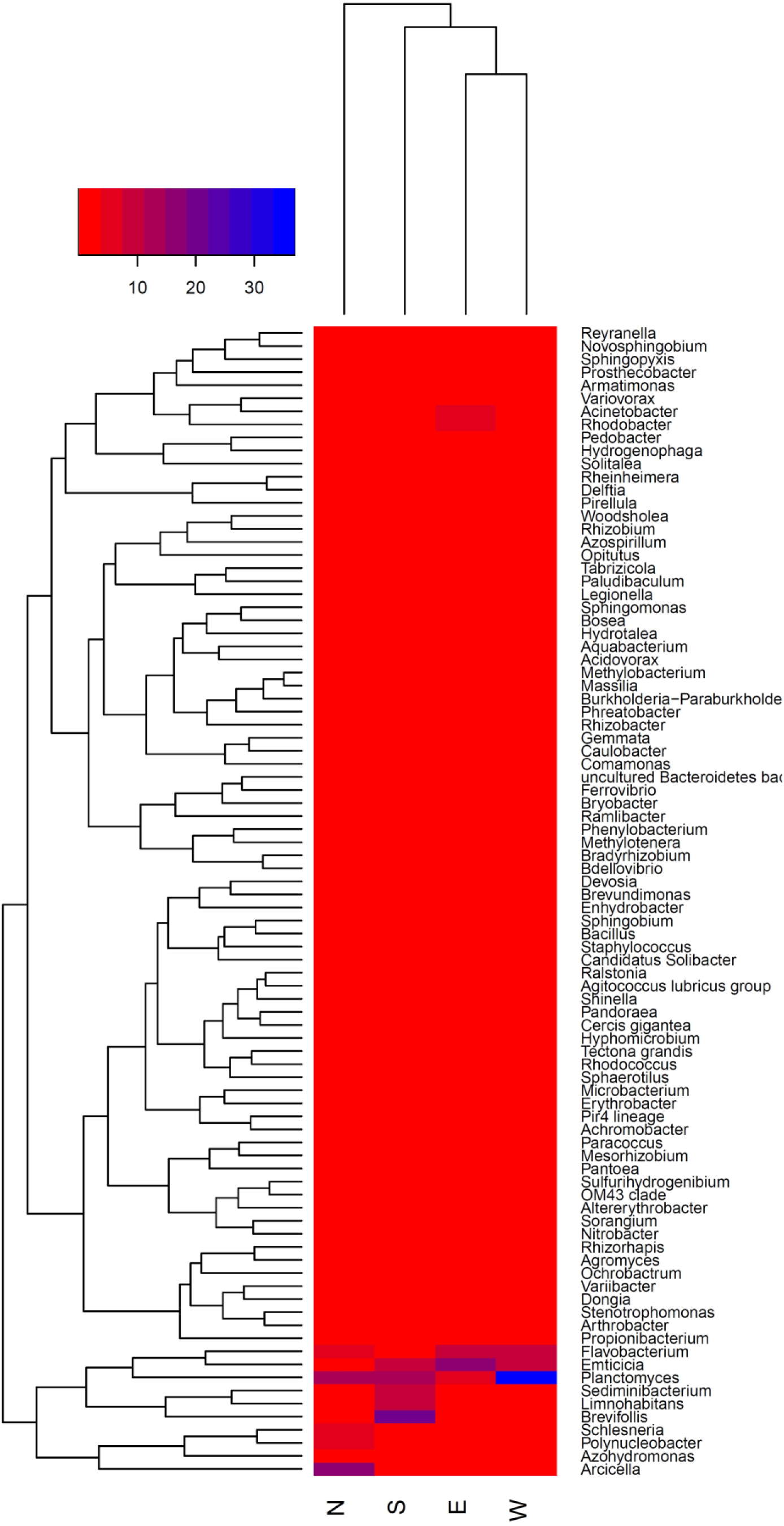
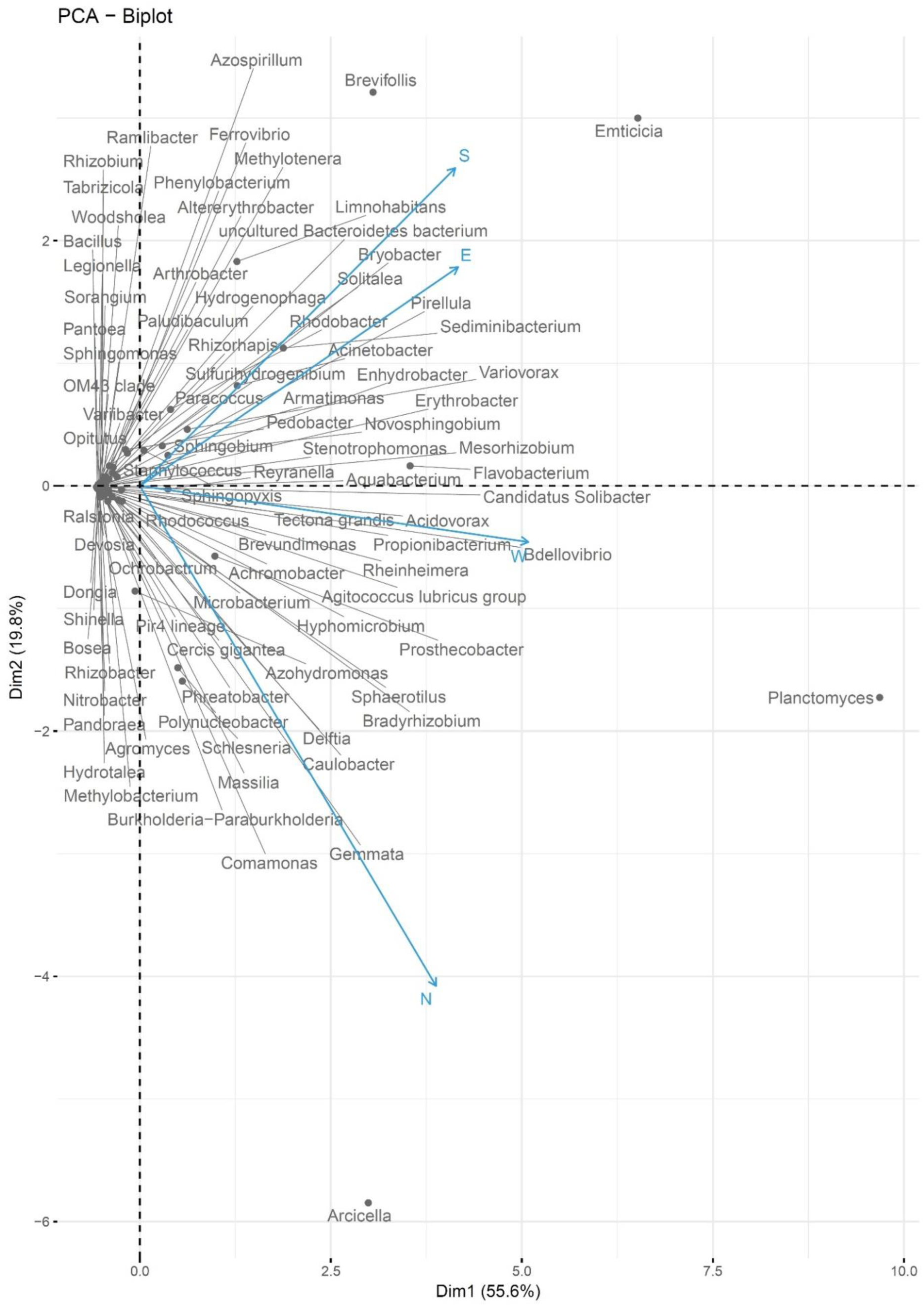
Heatmap drew with Bray Curtis distance method to analyze the comparative difference in microbial diversity among the metagenomic library of spring waters from east, west, south and north district (color key: red = lowest, blue = highest. **(b)** Principale component analysis of the microbial community of four districts (at genus level), the analysis showed a close relationship between the microbial community of east and south, while they are distantly related to the west, but north has totally different community structure forming an outgroup in the plot.

### 3.5 The relationship between physicochemical characteristics and microbial diversity

Multivariate canonical correspondence analysis of seven physicochemical parameter and seven dominant microorganisms of four districts showed a significant relationship. The distribution of dominant genus i.e. *Emticicia* and *Flavobacterium* in the east district was found to be closely dependent on the concentration of sodium (41.5ng/l) and calcium (35.2ng/l). Correspondingly, the relative abundance of the dominant genus of west district i.e. *Planctomyces* was found to be dependent on the concentration of Barium (9.761), Iron (10.823) and it was also influenced by the temperature. The dominance of *Arcicella* and *Schlesneria* in the North district was correlated with electroconductivity of water. Genus *Brevifolis* and *Sediminibactrium* in South district were correlated with the total dissolved solids present in the springs of South district **(Fig. 7)**.

**Fig. 7:**
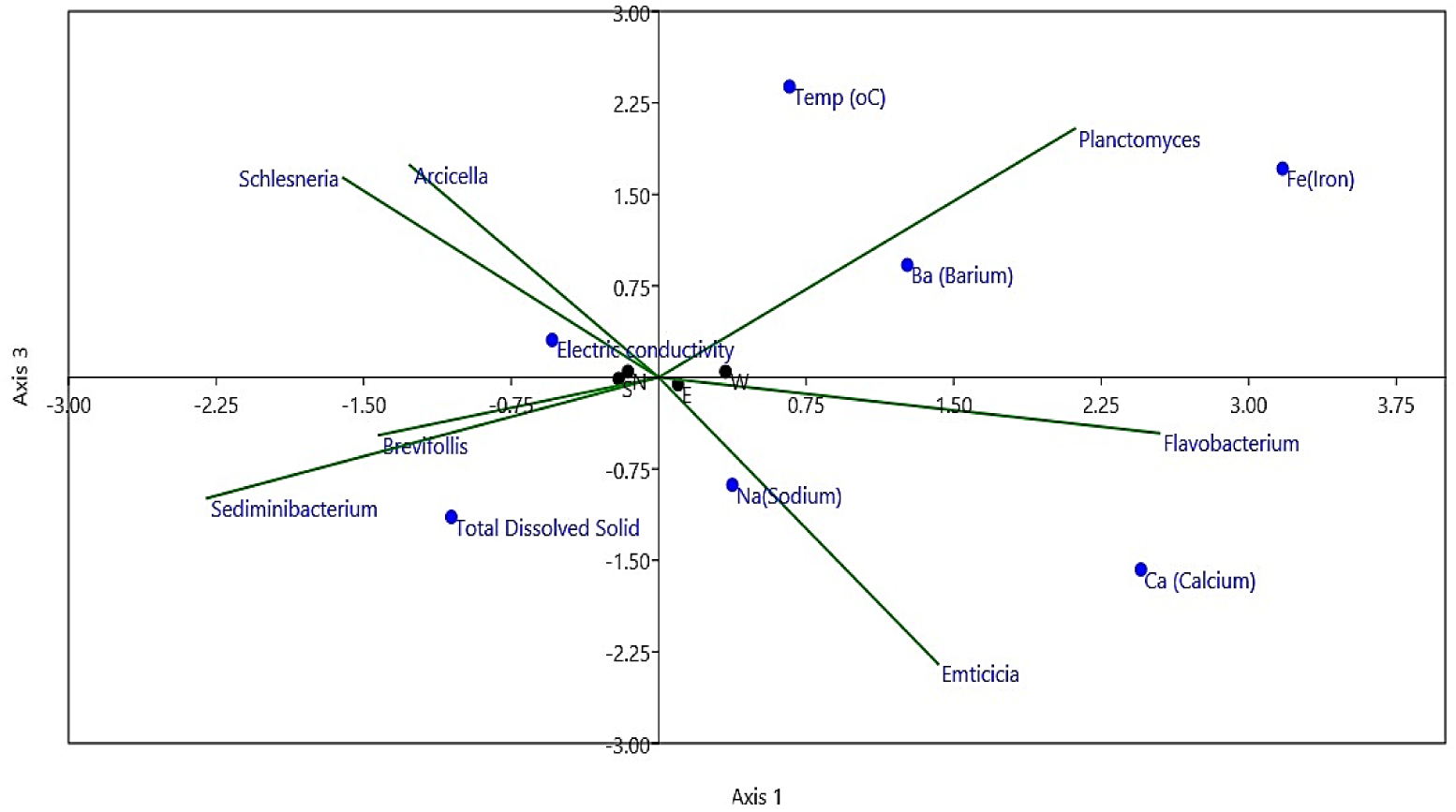
CCA plot showing the relationship between physicochemical parameters and microbial diversity (genus).

## 4 Discussion

Sikkim is a northeastern state lying in the lower region of Eastern Himalaya and neighbors to the countries China, Bhutan and Nepal. It is among the three hot spot ecoregions of India (The Western Ghat, Eastern Himalaya and Indo-Burma) with diverse fauna and flora. In topology, it has mountain terrain with wide altitude variation from 800-8000 meters. With a population of 6, 10, 577 it is the second smallest state of India. Majority of the population living in different altitudes are dependent on the natural spring waters for drinking and household purposes. Quality of life increases when people have access to safe drinking water with adequate sanitation. Better management of water resources to reduce different water-borne diseases and to make water safe for both potable and recreational purpose can save many lives. Water safety and quality are fundamental to human development and well-being (WHO, 2018). However, till date, no such studies have been conducted in Sikkim to determine the water structure of natural springs from microbiological or chemical aspects. Microbiological analysis is one the main facet of water quality measurement. Taxonomic profiling of dominant microflora can be an environmental indicator of water quality and can forecast future health threats in the surroundings. Knowing the ecology could be helpful in determining future water treatment protocols. But, limitation of culture-dependent methods has always been a barrier for comprehensive microbial ecology analysis. Development of metagenomics approaches and next-generation sequencing has allowed the scientist to overcome the barrier to determine the total microbial biodiversity of an environment. In this study, microbial ecology of the spring waters of Sikkim was determined by next-generation sequencing using variable regions of 16S rRNA gene (V3-V4). To our knowledge, this is the first report on the microbial ecology of spring waters of Eastern and Western Himalayan Region of India.

The study showed a major dominance of *Bacteroidetes*, *Proteobacteria*, *Verrucomicrobia* and *Planctomycetes* in all the springs of East (E), West (W), South (S) and North (N) districts **(Fig. 3a)**. Springs of East district was dominated by *Proteobacteria (41%)* while springs of West district was dominated by *Planctomycetes* (38.46%). *Planctomycetes* (7.54%) was least dominated in the East as compared to the West district (38.46%). *Planctomycetes* are a phylum of aquatic bacteria, with its habitat in fresh, marine and brackish water (Fieseler *et al*., 2004). They possess unusual characteristics such as intracellular compartmentalization and they lack peptidoglycan in their cell walls. Remarkably, few genera of this group like *Gemmata* even contain membrane-bound nucleoid similar to the eukaryotic nucleus (Boedeker *et al*., 2017). Although a recent study reported its close similarity to Gram-negative bacteria however in-depth genetic studies is still lacking (Boedeker *et al*., 2017). A unique bacterial phylum *Armatimonadetes* (8.08%) was recorded from the samples of the west districts which are not found in springs of other districts. Prior to official classification of *Armatimonadetes* by Tamaki *et al*., (2011), it was classified as candidate division of OP10, first identified by Hugenholtz *et al*., (1998) from Obsidian Pool, Yellowstone National Park (Hugenholtz *et al*., 1998; Tamaki *et al*., 2011; Lee *et al*., 2014). *Armatimonadetes* as a gram-negative oligotrophic aerobic bacterium was described by Lee *et al*., 2014. Spring water from South district was dominated by *Verrucomicrobia* (25.46%). *Verrucomicrobia* have only a few described species so far, most of the genus of the phylum is non-culturable and they are ubiquitous in freshwater and soil (Gupta, 2016; Griffiths and Gupta, 2007). The spring water of both North and East district was dominated by *Proteobacteria* (35.80 %) and *Bacteroidetes* (34.90%). The principal component analysis of relative phyla diversity also showed a close correlation between samples of the East and North districts while samples of South and West districts were distantly related (**Fig. 3b**).

The spring waters from four districts of Sikkim showed considerable differences among their dominant genus **(Fig. 5)**. The spring water of North was most diverse out of the three districts having major dominance of *Arcicella, Planctomycetes, Polynucleobacter, Schlesneria* and *Azohydromonas*. This difference in diversity can be due to the variance in the physicochemical parameter. North district has a comparatively lower temperature (17-22 °C) in comparison to other districts. *Arcicella* is aerobic gram-negative bacteria was first proposed by Nikitin *et al*., (2004) (Nikitin *et al*., 2004). Some of the novel species of *Arcicella* are isolated and identified from aquatic environment like *Arcicella aurantica* from stream water in Southern Taiwan (Sheu *et al*., 2018), *Arcicella rosea* from tap water (Kampfer *et al*., 2009) and *Arcicella rigui* from Niao-Song Wetland Park in Taiwan (Chen *et al*., 2013). Though the presence of *Planctomycetes* was found in all the four districts with lower relative abundance, its dominance was maximum at West (38.46%) and least at East district. The springs of East district was mainly dominated by the bacteria *Emticicia* (14.83%). *Emticicia* is a gram-negative bacterial genus belongs to family *Cytophagaceae* and they are ubiquitous in the aquatic environment (Schultz *et al*., 2013; Seo *et al*., 2015; Nam *et al*., 2016). A number of significant members from the genera were identified in aquatic systems *viz. Emticicia aquatica* (Seo *et al*., 2015), *Emticicia aquatilis* (Ngo *et al*., 2017), and *Emticicia fontis* (Nam *et al*., 2016). The bacterial genera *Flavobacterium* was found in all the other three districts except in the south. *Flavobacterium* a gram-negative bacterium belongs to phylum *Bacteroidetes* and are widely distributed in a freshwater ecosystem (Fernández-Gómez *et al*., 2013). *Flavobacterium* is responsible for bacterial cold water and bacterial gill diseases in different fish species (Strepparava *et al*., 2014). They are also opportunistic human pathogens, there are several reports on their association with pneumonia and bloodstream infection (Manfredi *et al*., 1999; Holmes *et al*., 1984). *Rhodobacter* and *Acinetobacter* were found in the south district which was absent in spring waters of other districts. The diversity of *Rhodobacter* in an aquatic system is ubiquitous; they are photosynthetic bacteria belonging to phylum *Proteobacteria* (IMHOFF *et al*., 1984). Some of the important *Rhodobacter sp*. identified from aquatic environment are *Rhodobacter adriaticus* isolated from Adriatic Sea (IMHOFF *et al*., 1984) ; *Rhodovulum aestuarii* isolated from brackish water collected from an estuary (Ramana *et al*., 2016); *Rhodobacter azollae* and *Rhodobacter lacus* isolated from different pond samples of Kukatpally, India (Suresh *et al*., 2017); *Rhodobacter vinaykumarii* a marine phototrophic *alpha-proteobacterium* from tidal waters in Visakhapatnam, on the east coast of India (Srinivas *et al*., 2007). There are also reports from Himalayan regions *viz. Rhodobacter changlensis* which was isolated from snow sample of Changlapass in the Indian Himalaya (Anil Kumar *et al*., 2007). Presence of *Acinetobacter* in rural drinking water systems dates back to 1989 (Bifulco *et al*., 1989). Since then, there are several reports of its presence in drinking water sources (Towner, 2006; Krizova *et al*., 2015; Radolfova-Krizova *et al*., 2016) and transmission of drug-resistant *Acientobacter* through oral route (Umezawa *et al*., 2015). Principle component analysis **(Fig. 6b)** of the microbial diversity of four districts showed East and South is correlated in genus wised distribution and they are distantly related with the Wast district but North district produced an out-grouped showing the divergence in diversity.

Canonical Correspondence Analysis **(Fig. 7)** confirmed the correlation between hydrochemistry and diversity. The diversity in the springs of the four districts is influenced by the concentration of different metallic compounds like sodium, calcium, barium, and iron. The diversity of *Emticicia* and *Flavobacterium* are influenced by the concentration of sodium and calcium while the diversity of *Planctomyces* was found to be dependent on the concentration of Barium and Iron. Along with the different chemical parameters, physical parameters like temperature, pH and electro-conductivity also had an influence on the diversity of microorganisms. The dominance of *Arcicella*, *Schlesneria*, *Brevifolis* and *Sediminibactrium* are influenced by the electroconductivity and total dissolved solids respectively. The results of this study significantly expand the current understanding of microbiology of the spring water of Sikkim and it also reports the comprehensive knowledge on microbial community structure which will help in near future to determine or to design any water treatment protocols or policies. This study also provides a brief insight of the physicochemical parameters of the spring water of Sikkim and their cross association with the indigenous microbial diversity.

## Acknowledgement

The authors wish to thank the State Institute of Rural Development Department, Government of Sikkim for their helping hand in providing information about springs locations and water sample collection. The authors also like to thank all the faculty member, non-teaching staff of the Department of Microbiology, Sikkim University for their continuous support and help throughout the study.

## Supplementary Files

**Table S1.** The quantity of gDNA as determined by Qubit Fluorometer and mean library fragment size.

| Sl. No. | Sample ID | The quantity of gDNA ng/<br>μl | Mean Library Fragment Size<br>(bp) |
| --- | --- | --- | --- |
| 1. | E | 31.7 | 605 |
| 2. | W | 15.0 | 539 |
| 3. | S | 19.8 | 598 |
| 4. | N | 18.5 | 605 |

**Table S2:** FastQ read statistics.

| Sample | Number of Reads | Total Bases | Observed OTUs |
| --- | --- | --- | --- |
| E | 248, 903 | 116,834,615 | 687 |
| W | 316,594 | 151,955,051 | 747 |
| S | 198,909 | 94,297,897 | 534 |
| N | 287,074 | 198,132,358 | 729 |

**Table S3:** Diversity Index

|  | S | W | N | E |
| --- | --- | --- | --- | --- |
| **Simpson_1-D** | 0.7499 | 0.7356 | 0.6988 | 0.6928 |
| **Shannon_H** | 1.534 | 1.49 | 1.36 | 1.368 |
| **Fisher_alpha** | 8.719 | 10.7 | 13.71 | 8.107 |
| **Chao-1** | 22 | 25 | 29 | 21 |

